# Co^2+^-mediated adsorption facilitates atomic force microscopy of DNA molecules at double-helix resolution

**DOI:** 10.1101/2025.05.29.655787

**Authors:** Mark Pailing, Taiana Maia de Oliveira, Maria M. Flocco, Bart W. Hoogenboom

## Abstract

Atomic force microscopy (AFM) has demonstrated the ability to resolve single DNA molecules in liquid at a spatial resolution that is sufficient to visualize the double helix structure and variations therein. Such variations can be due to inherent configurational flexibility and may be related to, e.g., DNA sequence, ionic screening, supercoiling, or protein binding. These AFM experiments require DNA to be adhered to a solid and preferably flat support. For high-resolution, in-liquid AFM studies so far, such adhesion has commonly been achieved using Ni^2+^ ions to electrostatically bridge between the negatively charged DNA and a negatively charged, atomically flat mica surface, yet Ni^2+^ ions tend to cause precipitation of salts on the surface, increasing the risk of AFM tip contamination and increasing the corrugation of the support surface, making it harder to distinguish secondary DNA structure. Here, we report on a sample preparation protocol that, instead, relies on Co^2+^ ions to adhere DNA to mica. While the Co^2+^ is similarly effective as Ni^2+^ for facilitating DNA adsorption onto mica, it leads to significantly reduced salt precipitation with the potential to provide enhanced reproducibility in high-resolution DNA imaging by AFM. We expect this to substantially facilitate high-resolution AFM studies of DNA in aqueous solutions.

## Introduction

Arguably no biological structure is as iconic as the DNA structure, originally discovered using X-ray crystallography (Watson and Crick, 1953). Since then, countless DNA structures have been solved using X-crystallography, showing B, A and Z-DNA along with many other non-canonical DNA forms such as triplex DNA and DNA quadruplexes (Egli, 2004; Wang et al, 1979; Rhee et al, 1999; Parkinson et al, 2002). However, such structure determination depends on crystallisation of short DNA segments and does not reflect the configurational flexibility of longer DNA molecules (≳15 base pairs) as present in the cell. Cryo-electron microscopy has facilitated structure determination of some more complex DNA arrangements, yet still depends on ensemble averaging of large numbers of identical molecules to achieve sub-nanometre resolution (Kato et al, 2009).

Atomic force microscopy (AFM) represents a complementary tool for imaging DNA molecules in a physiologically relevant environment that does not rely on averaging large numbers of molecules to obtain structural information. AFM has been used since the 1990s to study DNA structure, supercoiling and non-canonical forms of DNA (Hansma and Laney, 1996; Main et al, 2021). In addition, the development of high speed-AFM has allowed for the study of DNA conformational flexibility and dynamics (Yang et al, 2015; Kobayashi et al, 2007; Crampton et al, 2007). One of the appealing features of AFM is its ability to resolve DNA structures in physiologically relevant buffers (Thomson et al, 1996). As early as the mid-1990s, AFM provided measurements of the helical pitch at the level of individual molecules (Mou et al, 1995); and over the past decade, numerous AFM studies have achieved spatial resolution that has been high enough to identify the two strands of the double helix, e.g., showing variability of the helical pitch, revealing sequence-dependant structural deviations of canonical B-DNA, showing the structure of triplex forming DNA (Pyne et al, 2021) and more generally providing high-resolution images of DNA-protein interactions (Leung et al, 2012; Pyne et al, 2014; Pyne and Hoogenboom, 2016; Hamon et al, 2007; Kuchuk and Sivan, 2018; Ido et al, 2018; Kuchuk et al, 2019; Rhee et al, 1999).

However, for AFM to achieve such resolution, it is essentially to deposit DNA molecules on a solid substate. The most commonly used substrate is muscovite mica, which is formed from thin aluminium silicate layers held together by interstitial potassium ions (Tang et al, 2014; Hansma and Laney, 1996; Billingsley et al, 2014). Following cleavage between these layers, the exposed mica surface is atomically flat, facilitating AFM analyses in aqueous solution and exposing a honeycomb lattice with cavities at the sites of potassium ions, and a negative charge density of -1.0 to -1.7 mC/m^2^ (Pastré et al, 2003; Kan et al, 2015; Sushko, Shluger and Rivetti, 2006; Akpinar et al; 2019). To overcome the electrostatic repulsion between the mica surface and the also negatively charged phosphodiester backbone of DNA (at neutral pH), and thereby to facilitate DNA adsorption, various functionalization methods have been developed. These include the use of AP-mica (Pastré et al, 2003), 3-aminopropyltriethoxysaline modified mica (Shlyakhtenko et al, 2012), silylated mica (Bezanilla et al, 1995), Poly-L-Lysine (PLL), and polyethylene glycol-_b_-poly L-lysine (PEG-_b_-PLL) co-block polymers (Akpinar et al, 2019), which all result in a positive surface charge at neutral pH. However, the most commonly used and arguably most straightforward surface functionalization is by the use of divalent counterions.

The use of different divalent counterions has been extensively explored in the preparation of dried DNA samples for in-air AFM analysis. It was quickly established that transition metal ions including Ni^2+^, Co^2+^ and Zn^2+^ allowed for efficient and strong adsorption of DNA molecules to mica (Cheng et al, 2006). All these cations have radii of ≤ 0.74 Å, fitting in the cavities left by desorbed potassium ions at the cleaved mica surface (Hansma and Laney, 1996). It has been suggested that strong interactions between these cations and the -OH groups at the mica surface are further facilitated by their unpaired d-orbital electrons (Hansma and Laney 1996; Billingsley et al 2014). DNA adsorption was shown to be concentration dependant, with cation concentrations below ∼1 mM unable to produce a large enough positive charge to allow adsorption, whereas cation concentrations above ∼10 mM lead to effective repulsion due to excess counterion accumulation both at the mica surface and at the DNA (Hansma and Laney, 1996; Hansma and Laney, 1993).

Another commonly used counterion is Mg^2+^, which yields sufficiently strong DNA binding to facilitate AFM imaging in air, but it lacks unpaired d-orbital electrons and as such ions remains in dynamic equilibrium with the mica surface in solution (Billingsley et al, 2014; Piétrement et al, 2003). This can yield weak absorption of DNA on mica in solution, yet the DNA remains highly mobile, which is an advantage for studying molecular dynamics e.g. by high-speed AFM (Suzuki et al, 2010; Kobayashi et al, 2007), but this mobility has thus far compromised the spatial resolution that could be achieved by AFM. Alongside divalent cations, biological polyamines such as putrescine^2+^ and spermidine^3+^ have been used to adsorb DNA to mica (Sukhanova et al, 2015). These ions cause DNA to only weakly bind mica, enabling the study DNA dynamics and restriction enzyme activity using AFM (Lee et al, 2015). However, the increased DNA dynamics on the surface typically limits the spatial resolution (Pastré et al, 2010). By contrast, DNA binding by Ni^2+^ ions facilitates a range of binding strengths; at Ni^2+^ concentrations of ∼1 mM, DNA appears to be loosely bound and can still undergo conformational changes (Pyne et al, 2021), whereas at concentrations as high as 50 mM, strong binding occurs, in some cases yielding AFM topographies that are consistent with the predicted appearance of individual phosphate groups at the DNA surface (Ido et al, 2013).

However, the use of Ni^2+^ has its own disadvantages in that it is known to form various stable crystals in solution, which may include nickel hydroxide, nickel oxide and nickel chloride (if the initial salt used was nickel chloride). This results in precipitates at the mica surface, particularly at higher Ni^2+^ concentrations and during experiments involving longer incubation/imaging times. In practice, this implies two main obstacles that affect the reproducibility of high-resolution DNA imaging by AFM. Firstly, these nickel precipitates compromise the otherwise atomically flat background against which DNA and protein structures can be distinguished; and secondly, these precipitates represent a risk of AFM tip contamination, compromising overall resolution (Hsueh et al, 2010; Lide, 2008; Piétrement et al, 2003; Hansma et al, 1993).

Ideally, such counterion-based DNA adsorption on mica should be tuneable by counterion concentration, result in strong binding at higher concentrations for high-resolution AFM imaging, and yield little to no precipitation of salts from the solution. Noting the importance of unpaired d-electrons for stable DNA-mica binding, we here explore the use of Co^2+^ to adsorb DNA to mica for high-resolution AFM experiments in aqueous solution. Both Ni^2+^ and Co^2+^ have ionic radii, 0.69 Å and 0.72 Å, small enough to fit in the cavities left by desorbed K^+^ ions (radius: 0.74 Å) at the mica cleavage planes. Compared with nickel, cobalt has an additional unpaired d-electron, yet is not as reactive as iron, its other neighbour in the period table. We find that mica treated with Co^2+^ allows efficient and strong DNA binding and leads to significantly less surface contamination than mica treated with Ni^2+^, across of a range of concentrations. This results in more consistent high-resolution DNA imaging, as illustrated by robust double-helix resolution imaging against a cleaner background. DNA adsorbed with Co^2+^ also shows helical dimensions in agreement with those obtained in crystallographic studies, with a helical pitch of 3.4 +/-0.2 nm.

## Methods and materials Materials

Cobalt chloride hexahydrate (Sigma; Cat no: C-2644) and nickel chloride hexahydrate (Sigma; Cat. No: N6136) were dissolved to a final concentration of 0.1 M in MiliQ water and buffered to a pH of 7.4 using

12.5 mM HEPES. Imaging buffer (12.5 mM NaCl, 12.5 mM HEPES) was produced and buffered to either pH 7.4 or 4.7. All solutions were filtered using a 0.2 µm Acrodisc Syringe Filter with Supor Membrane (Pall Corporation; Cat No: 4612) and stored at 4°C. 6 mm sample disks were punched from ruby mica sheets (Agar Scientific; Cat No: G250-1) and cleaved immediately prior to use (Pyne and Hoogenboom, 2016).

### DNA production

496 base pair (bp) DNA was PCR amplified from lambda DNA (ThermoFisher, Cat. No: SD0011) using the forward primer 5’-CGATGTGGTCTCACAGTTTGAGTTCTGGTTCTCG-3’ and the reverse primer 5’-GGAAGAGGTCTCTTAGCGGTCAGCTTTCCGTC-3’ purchased from Integrated DNA Technologies. PCR reactions were conducted using dNTPs (ThermoFisher, Cat No: AB0196) and Phusion High-Fidelity DNA Polymerase (New England Biolabs, Cat. No: M0530S). PCR products were purified using a QIAquick PCR purification kit (Qiagen, Cat. No: 28104) and stored at 4°C.

### Mica functionalisation and DNA deposition

For studies assessing the amount of DNA adsorbed the mica surface, 2 µL 10 nM 496 base pair DNA was incubated in 38 µL pH 7.4 imaging buffer with Co^2+^ or Ni^2+^, at a final cation concentration of 1, 5, 10 or 50 mM, on freshly cleaved mica for 30 minutes in a humidified environment prior to imaging on the AFM. Humidified environments were produced by placing paper towels soaked in MiliQ water into a closed petri dish.

For studies assessing surface contaminations over time, mica was cleaved and 40 µL 10 mM pH 7.4 Co^2+^ or Ni^2+^ in imaging buffer was added the mica. Discs were incubated in a humidified environment for 15, 30, 45 and 60 minutes prior to imaging.

For studies assessing surface contamination over time with DNA, mica was cleaved and 38 µL 10 mM pH 7.4 Co^2+^ or Ni^2+^ in imaging buffer was added to mica. 2 µL 10 nM 496 base pair DNA was added to the mica. Discs were incubated in a humidified environment for 20, 40, 60 and 80 minutes prior to imaging.

For studies assessing surface contamination with different counterion concentrations, solutions of 1, 5, and 10 mM Co^2+^ or Ni^2+^ was buffered to pH 7.4 or pH 4.7 in imaging buffer. For bare mica assessment studies, 1, 5 and 10 mM Mg^2+^ was buffered to pH 7.4 or pH 4.7 in imaging buffer. Mica was cleaved and 40 µL of each condition was deposited on the mica. Discs were incubated in a humidified environment for 30 minutes prior to imaging.

For studies assessing DNA conformational flexibility, 3 mm discs mica were cleaved and 2.75 µL of pH 7.4 0.25, 0.5 and 1 mM Co^2+^, or 1 mM Ni^2+^, in imaging buffer was added. 0.25 µL 10 nM 496 base pair DNA was added and incubated in a humidified environment for 5 minutes. Following this, the imaging chamber was then flooded with 1 mL imaging buffer plus divalent ion solution at the same concentration as the initial adsorption. Samples were incubated for a further 10 minutes prior to imaging.

### Atomic Force Microscopy

AFM imaging was conducted at room temperature using a FastScan Bio microscope (Bruker). Images were collected using PeakForce Tapping (PFT) mode. Force curves were collected using a PeakForce Tapping amplitude of 10 nm at a height of 20nm. Images were collected at a frequency of 8 kHz using FastScan-D-SS (Bruker) cantilevers (resonance frequency in liquid ∼110 kHz and nominal spring constant 0.25 Nm^-1^). Imaging speeds of 2.5-3.5 Hz (line rates) and PeakForce setpoints of 50-100 pN were used. All images were collected using 512 × 512 pixels. Images were processed in Gwyddion 2.6 (Neĉas and Klapetek, 2012) by mean plane subtraction, median line by line alignment, level data by facet points and a gaussian filter of 1 nm applied. For images containing DNA, a mask was applied by threshold to mask DNA. Image flattening was completed as above excluding the masked DNA.

High speed AFM imaging was conducted at room temperature using a JPK NanoRacer microscope (Bruker). Images were collected using high speed AC mode with USC-F1.2-K0.15 (Nanoworld) cantilevers; resonance frequency in liquid ∼1200 kHz and nominal spring constant 0.15 N/m. Imaging speeds of 300 lines/s and a setpoint of ∼7 nm was used. All images were collected using 512 × 512 pixels. Videos were processed using NanoLocz software (Heath, Mickelwaite and Storer, 2024). Images were loaded and flattened using the “iterative peaks” auto flattening function, filtered with a 1-pixel gaussian filter and aligned using the “auto-align” function to reduce image drift. Time courses of DNA molecules were exported as .tiff and .gif formats.

### DNA circularity analysis

Following imaging processing, tiff format time courses of DNA molecules were opened in ImageJ. Image properties (pixel size, time per frame) were assigned using the image properties function. DNA molecules were analysed using the TrackMate plugin (Ershov et al, 2022). DNA molecules were identified by threshold to produce an outline of the DNA. Tracks were filtered to remove unwanted features and identify the tracked DNA using the “TrackScheme” function. “Tracks” and “Spots” for individual DNAs were exported and the circularity of individual DNA molecules for each frame of the video was saved. Variance in circularity of individual DNA molecules over the time course was calculated using the “VAR.P” function on Excel. Variance in circularity for individual DNA molecules was plotted against the ion concentration.

### Surface Roughness quantification

Surface roughness was assessed by recording 1 µm x 1 µm square images of mica incubated with divalent ions species. For images containing DNA, 1 µm x 1 µm square images were obtained, DNA molecules were masked to exclude them from the roughness measurements. These images were processed as in section 2.3 and the statistical quantities function was used on Gwyddion 2.6 and RMS surface roughness was recorded.

### Statistical analysis

Statistical analysis was conducted in Origin Pro 2020. Mean values and standard deviation were calculated using the formulae function. Statistical significance was conducted using Two Sample t-tests.

## Results

### Ni^2+^ and Co^2+^ adsorb similar amounts of DNA to mica across a range of concentrations

We set out to assess the overall amounts of adsorbed DNA molecules in the presence of Co^2+^ ions, compared with an established protocol with Ni^2+^ (Haynes et al, 2022). DNA was adsorbed for 30 minutes on mica in the presence of different concentrations of Ni^2+^ and Co^2+^, prior to imaging in solution. Figure 1 shows that both Ni^2+^ and Co^2+^ can be used to adsorb DNA to mica in solution across a range of concentrations, with similar efficiency and with sufficient adhesion strength to facilitate subsequent AFM imaging in solution. The number of DNA molecules adsorbed to the surface increased from 1 mM to 10 mM for both Co^2+^ and Ni^2+^. The figure shows that at 50 mM Co^2+^, the number of DNA molecules adsorbed to the surface decreased (Fig. 1). Our findings agree with published data that the maximum adsorption of DNA molecules occurs in the range of 1-10 mM of divalent ions (Kuchuk et al, 2019).

**Figure 1.**
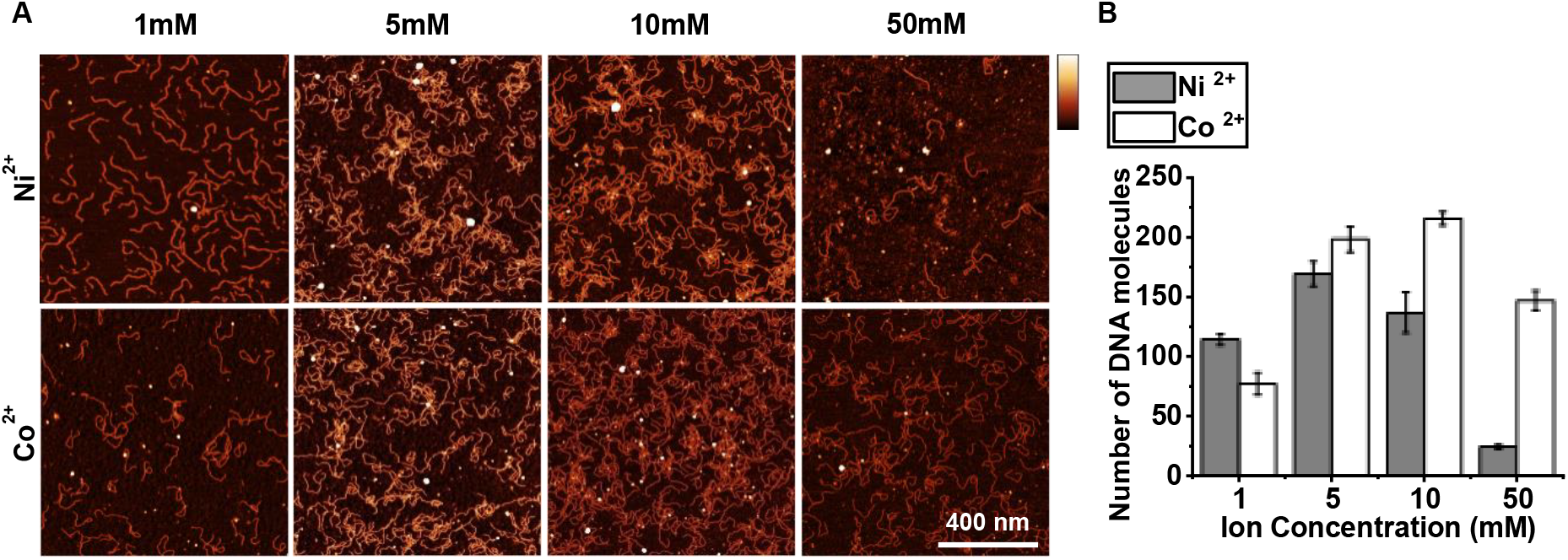
In liquid AFM imaging of 496 bp linear DNA adhered to mica in the presence of different concentrations of cobalt and nickel ions. A) 1 µm^2^ square images for DNA adsorbed to mica using increasing concentrations of Co^2+^ or Ni^2+^, in a buffer also containing 12.5 mM NaCl and 12.5 mM HEPES pH 7.4. (vertical scale: 5 nm). B) Quantification of the number of DNA molecules adsorbed to mica in images shown in A (n = 2 biological repeats). Error bars represent 1 standard deviation from the mean.

### Treatment of mica with Co^2+^ leads to less surface contamination than Ni^2+^ -

On closer inspection of the AFM images, we noted that the mica background appeared less corrugated in the Co^2+^ solutions than in the Ni^2+^ solutions, suggesting that cobalt salts precipitated less than nickel salts, thereby potentially avoiding a major downside of the use of Ni^2+^ in AFM experiments on DNA in solution. To test this, we exposed cleaved mica to solutions containing 10 mM Co^2+^ or Ni^2+^ in pH 7.4 imaging buffer, and recorded AFM images over a period of one hour. Mica incubated with Ni^2+^ shows a scatter of precipitates, visible as white dots in the images. Such precipitates are much less pronounced when Co^2+^ was used instead (Fig. 2A). More quantitatively, in Figure 2B, we found a marked increase in (RMS) surface roughness for Ni^2+^, from 0.28 nm at 15 minutes up to 0.46 nm at 60 minutes, at the resolution applicable in these images. By contrast, as recorded under identical conditions except for using Co^2+^ instead of Ni^2+^, the surface roughness remained at a baseline of ∼0.2 nm over 60 minutes when treated with Co^2+^. In brief, the presence of Co^2+^ ions lead to less precipitation of salts than is the case for Ni^2+^, with the difference more pronounced over longer incubation times.

**Figure 2.**
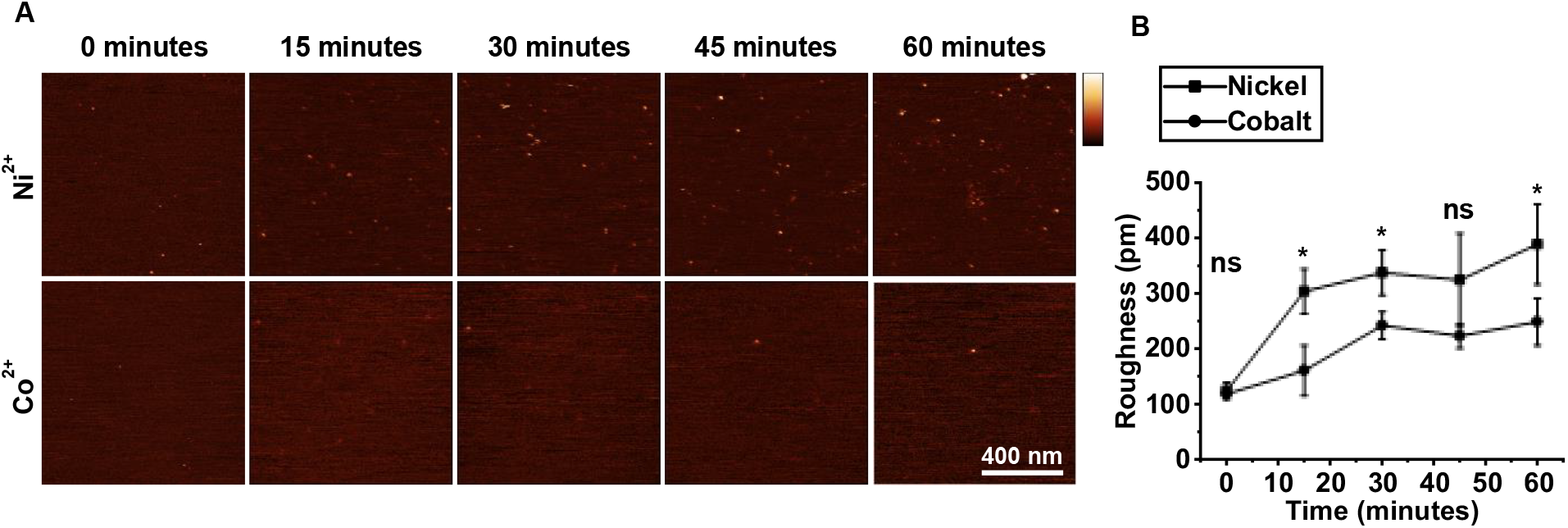
Surface roughness of mica incubated with Ni^2+^ and Co^2+^, as a function of time. A) 1 µm^2^ square AFM images recorded in 10 mM Co^2+^ or Ni^2+^, in imaging buffer, pH 7.4, as a function of incubation time (vertical scale: 5 nm). B) Quantification of the surface roughness of the images shown in A (n = 3 independent sample preparations). Error bars represent 1 standard deviation from the mean. T-tests were calculated * represents p < 0.05, ** represents p < 0.01, *** represents p < 0.001.

### Treatment of mica with Co^2+^ leads to less contamination than Ni^2+^ in the presence of DNA

Following our observation that Co^2+^ causes less mica contamination over time, we hypothesized that less contamination would develop in the presence of DNA, thereby enabling longer and clearer imaging conditions. To test this, mica was treated with 10 mM Co^2+^ or Ni^2+^ and DNA was incubated for 20 minutes. Images were taken every 20 minutes after the initial cleave. Higher surface contamination in Ni^2+^ treated mica is shown by large circular precipitates taller than 2 nm present throughout all images (Fig. 3A). Smaller deposits smaller than 2 nm are also present throughout the time course and increase in density. Smaller contamination is also present throughout the time course when treated with Co^2+^, however these are less dense than in Ni^2+^ images. Large deposits (> 2 nm) are present in Co^2+^ treated mica from 60 minutes onwards. To quantify surface contamination, a mask was applied to each image to remove DNA from roughness measurements. Roughness (RMS) was then acquired for the unmasked features. Roughness of mica treated with Co^2+^ is less contaminated following the initial 20-minute incubation to deposit the DNA when compared to Ni^2+^, and 80 minutes after initial treatment, the surface roughness is significantly lower when mica is treated with Co^2+^ Surface roughness increases at a slower rate compared to Ni^2+^ over the whole 80-minute time course.

**Figure 3.**
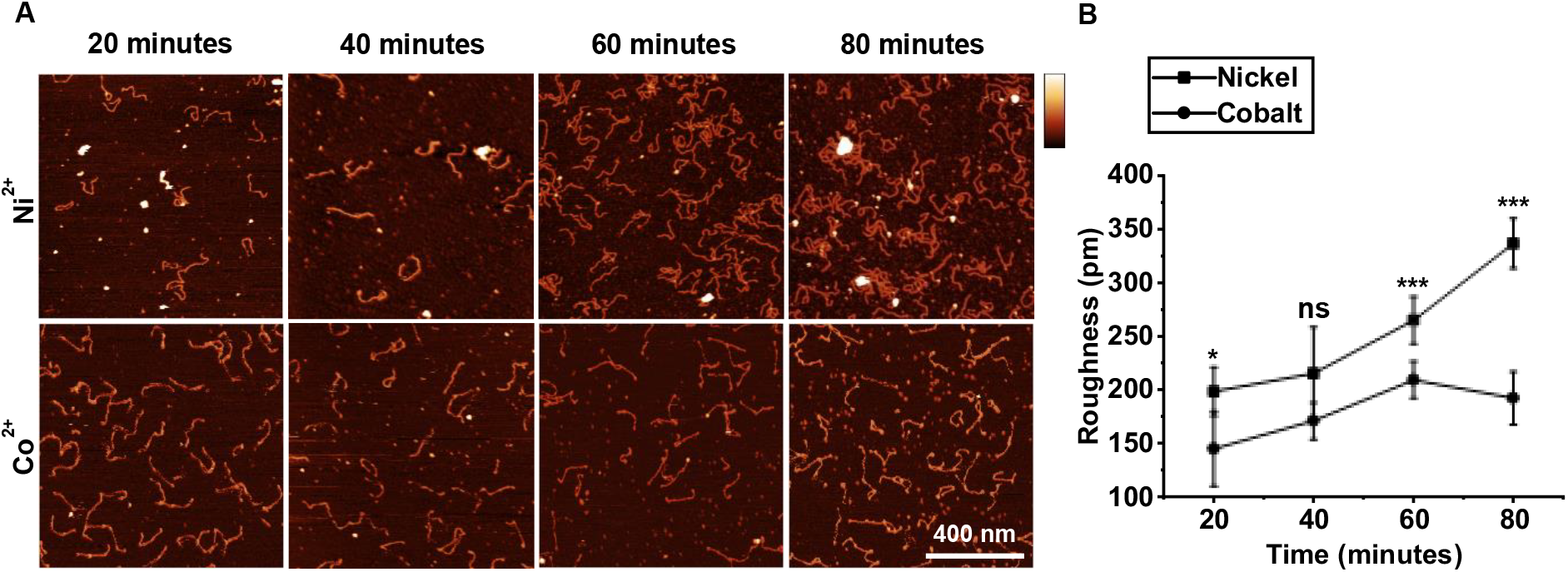
Surface roughness of mica treated with 10mM Co^2+^ or Ni^2+^ in the presence of DNA. A) AFM images of DNA adsorbed to mica treated with 10 mM Co^2+^ or Ni^2+^. B) Mean RMS roughness of mica treated with 10 mM Co^2+^ or Ni^2+^ in the presence of DNA (biological replicates n=4) (vertical scale = 5 nm). Error bars represent 1 standard deviation from the mean. T-tests were calculated * represents p < 0.05, ** represents p < 0.01, *** represents p < 0.001.

### Treatment of mica with Co^2+^ leads to less surface contamination than Ni^2+^ across a range of concentrations

To further quantify this observation, we measured the surface roughness as a function of Ni^2+^ and Co^2+^ concentration, compared with Mg^2+^ containing buffers as a non-transition metal control with identical Debye screening length. The next steps of our study was to assess surface contamination of mica associated with divalent ions across a range of concentrations and pH. Mica surface roughness of mica increases with Ni^2+^ and Co^2+^ concentration (Fig. 4A). Consistent with our previous findings, mica incubated with Co^2+^ has reduced surface roughness across the concentration range compared to Ni^2+^. Surface roughness is significantly reduced when mica is incubated with Co^2+^ compared with Ni^2+^ at both 1 mM and 10 mM concentrations (Fig. 4A and 4B). Interestingly, mica incubated with 1 mM Co^2+^ has a lower surface roughness than mica incubated with the Mg^2+^ control. Taken together these data clearly show that treatment of mica with Co^2+^ leads to a significant reduction in surface contamination than with when mica is treated with Ni^2+^ across a range of concentrations.

**Figure 4.**
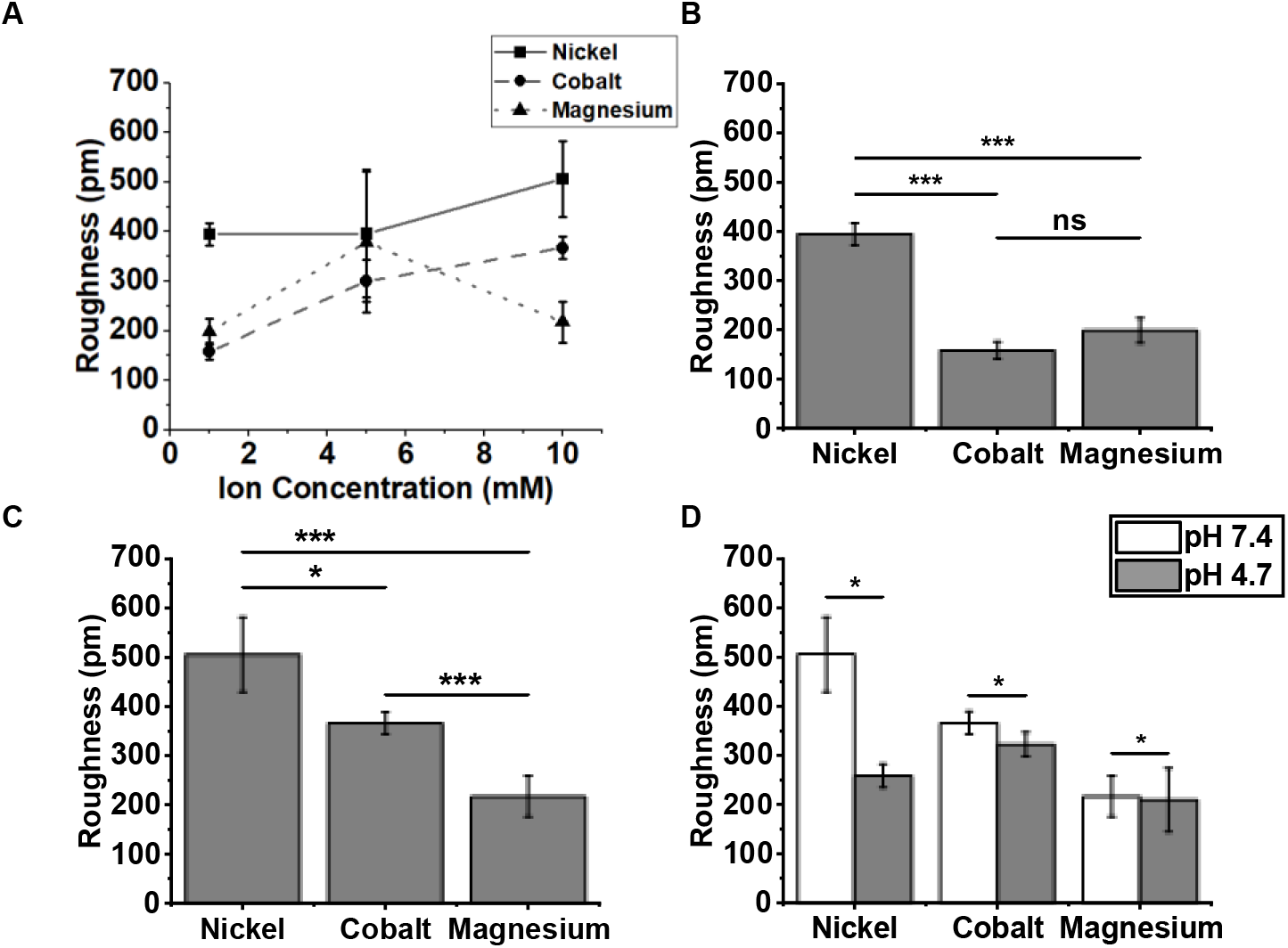
Surface roughness of mica incubated as a function of concentrations of divalent ions and of pH. A) Surface roughness of mica incubated with increasing concentrations of Co^2+^, Ni^2+^ or Mg^2+^ in pH 7.4 imaging buffer (n = 4 independent sample preparations). B) Surface roughness of mica incubated with 1 mM Co^2+^, Ni^2+^ or Mg^2+^ in pH 7.4 imaging buffer for 30 minutes (n = 4 independent sample preparations). C) Surface roughness of mica incubated with 10 mM Co^2+^, Ni^2+^ or Mg^2+^ in pH 7.4 imaging buffer for 30 minutes (n = 4 independent sample preparations). D) Surface Roughness of mica incubated with 10mM Co^2+^, Ni^2+^ or Mg^2+^ in pH 7.4 or pH 4.7 imaging buffer for 30 minutes (n = 4 independent sample preparations). Error bars represent 1 standard deviation from the mean. T-tests were calculated * represents P < 0.05, ** represents P <0.01, *** represents P < 0.001.

The observed precipitation in Ni^2+^-containing solutions has previously been attributed to nickel hydroxide, nickel chloride and/or nickel oxide (Billingsley et al, 2014; Hsueh et al, 2010). Our comparison with Co^2+^ provides a complementary perspective to the nature of the Ni^2+^ induced precipitation. Cobalt chloride exhibits a lower solubility in water than nickel chloride, 53 g/100 ml (Dean, 1987,p.2-12) compared to 64.2 g/100 ml (Prager, 1995, p.868), such that one would predict more precipitation of chloride-containing crystals for Co^2+^ than for Ni^2+^, which appears to contradict our experimental observation that there is less precipitation for Co^2+^. However, the bulk solubility of nickel hydroxide is less than half than that of cobalt hydroxide (1.5 mg/L compared to 3.2 mg/L) (Lide, 2009), suggesting that hydroxide may be the main responsible for the observed precipitation in the presence of Ni^2+^. To test this hypothesis, we reduced the concentration of OH^-^in solution by reducing the pH from 7.4 to 4.7 and measured the surface roughness as before. As expected for roughness due to precipitation of nickel hydroxide, the measurements at lower pH showed a significant and two-fold reduction in RMS roughness, Instead, for Co^2+^, the differences between the roughness for the two pH values were smaller than the standard deviations of the roughness measurements (Fig 4D). We therefore suggest that the observed precipitations may be attributed to nickel hydroxide and explain the better performance of Co^2+^ in this respect – at least in part – by the better solubility of cobalt hydroxide compared with nickel hydroxide.

### DNA conformational flexibility can be studied through fine tuning Co^2+^ concentration

We assessed whether the adsorption strength could be altered to study DNA conformational flexibility by fine tuning the concentration of divalent ions in solution (Lee et al, 2015). To this end, DNA was adsorbed to mica treated with various concentrations of divalent ions, followed by flooding the sample chamber with buffers containing the same species of ion at the same concentration as in the initial DNA adsorption. At higher Co^2+^ concentrations (1 mM), DNA molecules were relatively stably bound, with only the ends of the DNA molecules showing some conformational flexibility (Supplementary Videos 1- 6). Figure 5A shows a sequence of images of DNA adsorbed to mica at a lower, 0.5 mM Co^2+^ concentration. As a guide to the eye, the outline of the DNA in the first frame was traced and added to all subsequent images as a white dashed line. This sequence illustrates DNA mobility on the surface at a low divalent cation concentration; similar changes in conformation were found for numerous DNA molecules under such conditions (Supplementary Videos 7-10). For 0.25 mM Co^2+^, the DNA showed rather variable behaviour. Some DNAs were stably adsorbed and only flexible at the ends, whereas others were flexible across the whole molecule (Supplementary Videos 11-18), in some cases apparently affected by the movement of the AFM tip as it scanned the molecule (Supplementary Videos 14 and 16). The variability observed suggests that at lower Co^2+^ concentrations, Co^2+^ integration in mica is less uniform, producing areas with higher positive charge and thus stronger DNA adsorption. For comparison, DNA was also adsorbed used 1 mM Ni^2+^, which resulted in DNA behaviour similar to that of DNA molecules adsorbed with 0.25 mM Co^2+^ (Supplementary Videos 19-26).

**Figure 5.**
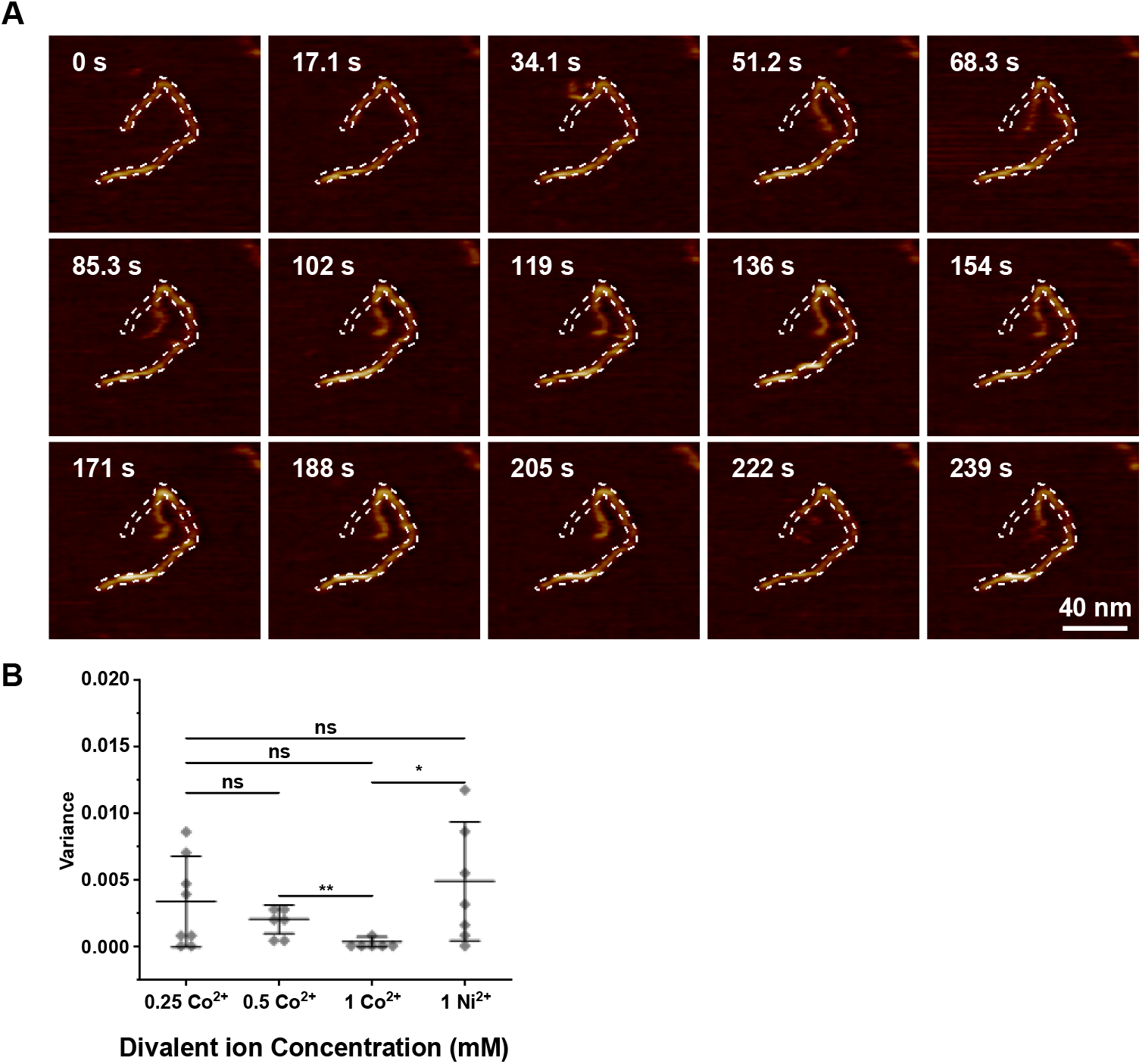
Higher Co^2+^ concentration results in stronger DNA adsorption and reduced conformational flexibility. A) AFM images of DNA adsorbed to mica treated with 0.5 mM Co^2+^ (vertical scale = 1.25 nm). B) Variance in circularity of individual DNAs adsorbed to mica with different divalent ions, ionic species and concentration indicated in figure (biological replicates n = 3). Error bars represent 1 standard deviation from the mean. T-tests were calculated * represents P < 0.05, ** represents P <0.01, *** represents P < 0.001.

For a more quantitative representation of these results, the outlines of individual DNA molecules were traced using threshold detection and the circularity of the DNA outline was recorded as a function of time (Ershov et al, 2022). The variance in this circularity was used as a measure of DNA flexibility, with higher flexibility implying a weaker DNA binding to the mica substrate. Figure 5B shows the results of this quantification for different Co^2+^ concentrations and for 1 mM Ni^2+^. The trends in Figure 5B confirm the more qualitative observations described above. Moreover, the lower flexibility 1 mM Co^2+^, compared with 1 mM Ni^2+^, is consistent with stronger binding and thereby with the larger amount of DNA on the mica substrate (for Co^2+^ vs. Ni^2+^ at 1 mM) as in the AFM imaging in Figure 1.

### High resolution imaging of DNA secondary structure of DNA adsorbed via Co^2+^

Finally, we assessed to what extent, aided by the reduced salt precipitation and stable DNA adsorption, the cobalt-mediated DNA adsorption facilitated imaging of DNA at sufficient spatial resolution to distinguish the major and minor grooves of the double helix (Fig. 6A,B). The typical double-banded structure of the double helix, somewhat widened by tip convolution, could clearly be resolution as well as highlighting the stability of imaging DNAs adsorbed to mica via Co^2+^ (Pyne et al, 2014). Fig. 6B show in detail the major and minor groove of the right-handed helix that is synonymous with B-DNA. The helical pitch (Fig. 6C) was measured across 3 samples and estimated to be 3.37 ± 0.18 nm, consistent to within the error with the pitch length in crystallographic studies (Watson and Crick, 1953). Finally, the height of the DNA (Fig. 6D), subject to only minor compression due to the AFM tip, was close to the ∼2.0 nm DNA diameter expected based on in crystallographic studies (Pyne et al, 2014; Watson and Crick, 1953).

**Figure 6.**
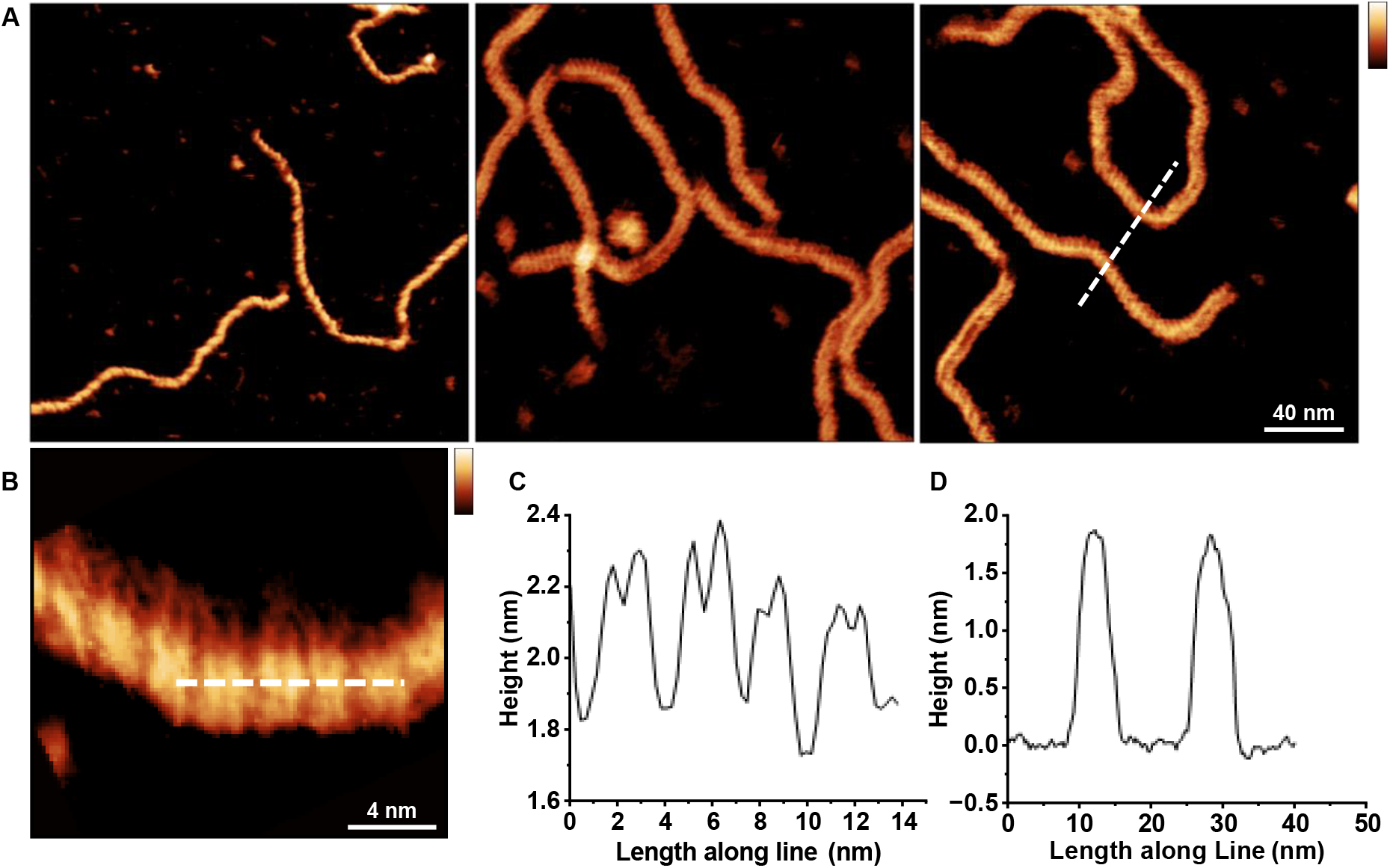
Double-helix resolution imaging of DNA adsorbed to mica via Co^2+^. A) High resolution AFM images of the DNA double helix adsorbed to mica in a buffer containing 10 mM Co^2+^ (vertical scale = 4 nm). B) Higher-magnification view of DNA, showing the double-banded structure representative of the DNA double helix. (vertical scale = 2 nm). C) Height profile along the DNA, as per white dashed line in B averaged over a width of 5 nm. D) Height profile across two DNA molecules, as per the white dashed line in A, averaged over a width of 5 nm.

## Discussion

Cleaved mica is the substrate of choice for AFM imaging due to its atomically flat surface (Hansma and Laney, 1996). However, the cleaved mica surface in water has a net negative charge and therefore repels the negatively charge backbone of DNA. This repulsion can be overcome by divalent cations, facilitating DNA adsorption to mica by providing a positive charge that bind both mica and DNA (Pastré et al, 2003). The properties of various divalent ions for DNA adsorption to mica have been extensively studied using AFM in air and liquid (Hansma and Laney, 1996; Billingsley et al, 2014; Cheng et al, 2006; Heenan and Perkins, 2019; Hseuh et al, 2010; Pastré et al, 2003). Smaller cations are able to fit into depressions in the mica that form when potassium ions embedded in the mica lattice are lost to solution.

Historically Ni^2+^ has been commonly used as the divalent ion for AFM imaging of DNA in solution. This is likely due to the ease of use, and Ni^2+^ is the smallest ion that facilitates stable DNA adsorption (Hansma and Laney 1996). However, Ni^2+^ is known to produce salt deposits on mica, which can impact imaging quality and stability (Hsueh et al, 2010; Main et al, 2021; Pyne and Hoogenboom, 2016). Here we have shown that Co^2+^ facilitates DNA binding to mica while producing less surface contamination than Ni^2+^ (Fig 4). Co^2+^ has a similar ionic radius as Ni^2+^ (0.72 and 0.69 respectively) (Hansma and Laney, 1996). Our data showed that Ni^2+^ surface contamination was pH dependant whereas Co^2+^ surface contamination was not (Fig 4). We therefore believe that Ni^2+^ surface contamination is caused by nickel hydroxide as OH-ions are reduced in lower pH buffers. This is line with previously published data which suggest that Nickel can form metal or metal-hydroxide layers on mica that are extremely sensitive to pH changes (Hseuh et al, 2010).

We also noted that double helix resolution imaging was readily achieved with Co^2+^ and produced DNA dimensions in line with previous AFM studies (Pyne et al, 2014) and crystallographic studies (Watson and Crick, 1953), possibly facilitated by the stronger binding of DNA to the mica substrate (at the same cation concentration) than for Ni^2+^. Current theories posit that unpaired d-orbital electrons are essential in forming contacts with mica and DNA. Co^2+^ has two unpaired d-orbital electrons whereas Ni^2+^ has one. The differing physical characteristics of Ni^2+^ and Co^2+^ (combined with reduced tendency for surface contamination) may be responsible for the stable high-resolution imaging of the DNA double helix at lower forces seen in our data. This extra unpaired d-orbital electron in Co^2+^ may also be responsible for the increased adsorption strength at lower concentrations compared to Ni^2+^ (Fig. 5). We have also shown that at lower concentrations, 1mM, Co^2+^ yields more DNA molecules adsorbed to the mica substrate than Ni^2+^ (Fig. 1).

While at the concentrations used, neither Co^2+^ nor Ni^2+^ can be considered physiological, Ni^2+^ is toxic to cells at lower concentrations and has been shown to cause cell death at concentrations above 250 µM in monocytes (Jakob et al, 2017) and osteoblasts (Navratilova et al, 2024). By comparison Co^2+^ is cytotoxic (at higher concentrations >400 µM, accumulating in the nucleus alongside chromatin suggesting a potential interaction with DNA or nuclear proteins (Ortega et al, 2009). This toxicity to cells does not noticeably affect the structure of individual DNA molecules though, given the good match between, on one hand, dimensions as measured by AFM here and elsewhere(Ido et al, 2013; Pyne et al, 2014), and on the other hand crystallographic studies (Watson and Crick, 1953).

We have here reported on the use of Co^2+^ as alternative divalent ion to facilitate DNA adsorption to mica and thereby high-resolution AFM imaging of DNA in solution. We have shown that Co^2+^ enables DNA adsorption to mica at lower concentrations than the (for in-solution AFM imaging) commonly used Ni^2+^. Furthermore, for the same cation concentrations, Co^2+^ led to reduced precipitation of salts at the mica surface compared to Ni^2+^. As is the case for Ni^2+^, Co^2+^ can be titrated to tune the DNA binding to the surface, thereby providing a means to optimize experimental conditions depending on the needs for imaging conformational changes (requiring only loose binding) or for highest-resolution imaging (benefitting from strong DNA binding to the surface). Such high-resolution imaging of Co^2+^-bound DNA on mica readily yielded images of sufficient quality to identify the major and minor grooves of the DNA double helix. Taken together, Co^2+^ represents an alternative divalent ion for AFM imaging of DNA that allows stable DNA imaging with, compared to the commonly used Ni^2+^, reduced risk of surface and AFM tip contamination in the form of precipitated salts.

## Notes

### Competing Interest Statement

B.W.H. holds an executive position at AFM manufacturer Nanosurf. Nanosurf did not play any role in the design or execution of this study.

### Funding

- Biotechnology and Biological Sciences Research Council: BB/W510026/1, BB/W019345/1
- Engineering and Physical Sciences Research Council: EP/M028100/1

